# High-sensitivity calcium biosensor on the mitochondrial surface reveals that IP3R channels participate in the reticular Ca^2+^ leak towards mitochondria

**DOI:** 10.1101/2022.12.02.518840

**Authors:** Yves Gouriou, Fabrice Gonnot, Ludovic Gomez, Gabriel Bidaux

## Abstract

Genetically encoded biosensors based on fluorescent proteins (FPs) are widely used to monitor dynamics and sub-cellular spatial distribution of calcium ion (Ca^2+^) fluxes and their role in intracellular signaling pathways. The development of different mutations in the Ca^2+^-sensitive elements of the cameleon probes has allowed sensitive range of Ca^2+^ measurements in almost all cellular compartments. Region of the endoplasmic reticulum (ER) tethered to mitochondria, named as the mitochondrial-associated membranes (MAMs), has received an extended attention since the last 5 years. Indeed, as MAMs are essential for calcium homeostasis and mitochondrial function, molecular tools have been developed to assess quantitatively Ca^2+^ levels in the MAMs. However, sensitivity of the first generation Ca^2+^ biosensors on the surface of the outer-mitochondrial membrane (OMM)do not allow to measure μM or sub-μM changes in Ca^2+^ concentration which prevents to measure the native activity (unstimulated exogenously) of endogenous channels. In this study, we assembled a new ratiometric highly sensitive Ca^2+^ biosensor expressed on the surface of the outer-mitochondrial membrane (OMM). It allows the detection of smaller differences than the previous biosensor in or at proximity of the MAMs. Noteworthy, we demonstrated that IP3-receptors have an endogenous activity which participate to the Ca^2+^ leak channel on the surface of the OMM during hypoxia or when SERCA activity is blocked.

## Introduction

Initially, the cytotoxic role of calcium ions (Ca^2+^) in ischemia was published over 40 years ago (Schanne et al., 1979; Schlaepfer and Bunge, 1973). The Ca^2+^ overload results from an unbalance of cell homeostatic pathways regulating Ca^2+^ influx, efflux and release from internal stores. Release of Ca^2+^ from the endoplasmic reticulum (ER) has been suggested to be the initial signal for ER dysfunction in ischemia (Paschen and Doutheil, 1999). The alteration of Ca^2+^ ATPase pumps due to the lack of energy supply, uncovers the preexisting ER calcium leakage through different channels and receptors participating in the ischemia-induced Ca^2+^ overload (Lemos et al., 2021). In non-excitable cells, the main Ca^2+^ release channel is the inositol 1,4,5-trisphosphate receptor (IP3Rs). IP3R channels are key elements of Ca^2+^ signaling machinery and reside in close proximity to the interface between ER and mitochondria microdomains to facilitate the transfer of Ca^2+^ ions (Gomez et al., 2016; Paillard et al., 2013; Szabadkai et al., 2006). In ischemic condition, the Ca^2+^-sensing receptor (CaR) has been shown to be activated in different models of ischemia/reperfusion (Lu et al., 2010; Paquot et al., 2017; Yan et al., 2010). These receptors elicit phospholipase C-mediated inositol triphosphate (IP3) formation, leading to a cytosolic Ca^2+^ elevation. Yet it remains unclear if IP3Rs could participate in both ER Ca^2+^ leak and cytosolic Ca^2+^ overload, not only at the early phase of the reoxygenation (Bruno et al., 2001) but also during the hypoxic period (Szydlowska and Tymianski, 2010). Thanks to the development of new biosensors this question can now be assessed by using targeted Ca^2+^-sensitive fluorescent proteins.

The engineering of genetically encoded fluorescent biosensors, based on green fluorescent protein (GFP) has expanded the versatility of metabolites quantification in signaling pathway networks study. Since its discovery in the 1990s (Tsien, 1998), GFP mutants have been extensively developed in a wide range of fluorescent proteins (FPs) with optimized brightness, photostability, folding and pH sensitivity. These optimizations of FPs have allowed the generation of robust tunable FRET-based biosensors to study Ca^2+^ signaling pathways. Indeed, the most common FRET biosensors are the Ca^2+^ sensitive cameleons based on CFP and YFP variants linked together by a Ca^2+^ binding domain from calmodulin and a calmodulin-binding domain from M13 skeletal-muscle myosin light-chain kinase (Palmer et al., 2006). Originally described in 2006 by Palmer et al., the Dcpv (cameleons) has been evolved into several variants: D1, D2, D3, D4 with different Ca^2+^ affinities and different cellular localization signals (Palmer et al., 2006). Cytosolic, mitochondrial and reticular cameleons Ca^2+^ biosensors have been generated and in 2010, Giacomello et al. have developed a GFP-based Ca^2+^ probe (N33D1cpv) localized on the outer mitochondrial membrane (OMM) suitable to monitor “Ca^2+^ hotspots” which means high Ca^2+^ levels from 1μM to 200-300μM (Giacomello et al., 2010) in a limited cellular area. The targeted sequence of the biosensor was based on the first 33 amino acids of TOM20 (N33), an endogenous protein of the OMM. Currently, there is no biosensor available to measure small variations of Ca^2+^ at the mitochondrial-associated membranes (MAMs) level.

In the present study, we assembled a new mitochondrial-surface GFP-based Ca^2+^ indicator derived from the GFP-based Ca^2+^ biosensor N33D1cpv, with a higher affinity for Ca^2+^ allowing sensitive Ca^2+^ measurements at the cytosolic surface of the OMM. By means of N33D3cpv biosensor, we showed that either genetic suppression or the pharmacological inhibition of endogenous IP3R activity reduced the speed of ER Ca^2+^ leak on the surface of the OMM during hypoxia.

## Results and Discussion

### Generation of a new mitochondrial-surface targeted GFP-based Ca^2+^ indicator

First of all, we substituted the D1 Ca^2+^-binding domain of N33D1cpv (Giacomello et al., 2010) by the D3 Ca^2+^-binding domain of the cytosolic biosensor D3cpv, which has a higher Ca^2+^ affinity. This new Ca^2+^ indicator was called N33D3cpv (Figure 1A). The D3 ligand domain has a dissociation constant (Kd) of 0.6μM that is particularly adapted to the range of low intensity Ca^2+^ variations in the cytosol (range of Ca^2+^ changes from 0.1μM to 10μM). By means of confocal imaging analysis, we validated the N33D3cpv localization around the outer-mitochondrial membrane (OMM) as described for N33D1cpv indicator (Giacomello et al., 2010). Cells transfected with N33D3cpv indicator together with a mitochondrial staining (mitotracker deep-red) showed a donut-like N33D3cpv fluorescence while mitotracker deep-red labelled the interior of mitochondria (Figure 1B, left panel). Confocal images using ERtracker red showed also that the probe is not only facing the ER (Figure 1B, right panel). Finally, we took advantage of the original protocol published by Palmer A. and Tsien R. (Palmer and Tsien, 2006), to perform an *in situ* calibration in order to compare with N33D1cpv the two key parameters, Ca^2+^ concentration and the dynamic range, for the N33D3cpv indicator (Figure 1C). We found a Kd of 1,5μM for N33D3cpv with the Ca^2+^ titration curve (Figure 1C) whereas N33D1cpv was reported having two Kd at: 18,61μM and 135,41μM (Giacomello et al., 2010). We also measured for each set of experiments the dynamic range for each probe. Briefly, Ca^2+^-buffered and Ca^2+^-saturated solutions were applied on permeabilized H9c2 rat cardiomyoblast cells, and variation of [Ca^2+^] outside OMM ([Ca^2+^]_OMM_) was measured after IP3-mediated Ca^2+^ release by ATP stimulation (Figure 1D). We observed that the variation of FRET ratio, triggered by ATP stimulation, was 6-fold greater in N33D3cpv compared to the original N33D1cpv. This confirms the higher sensitivity of N33D3cpv indicator for physiological Ca^2+^ fluxes on the surface of the OMM.

**Figure 1.**
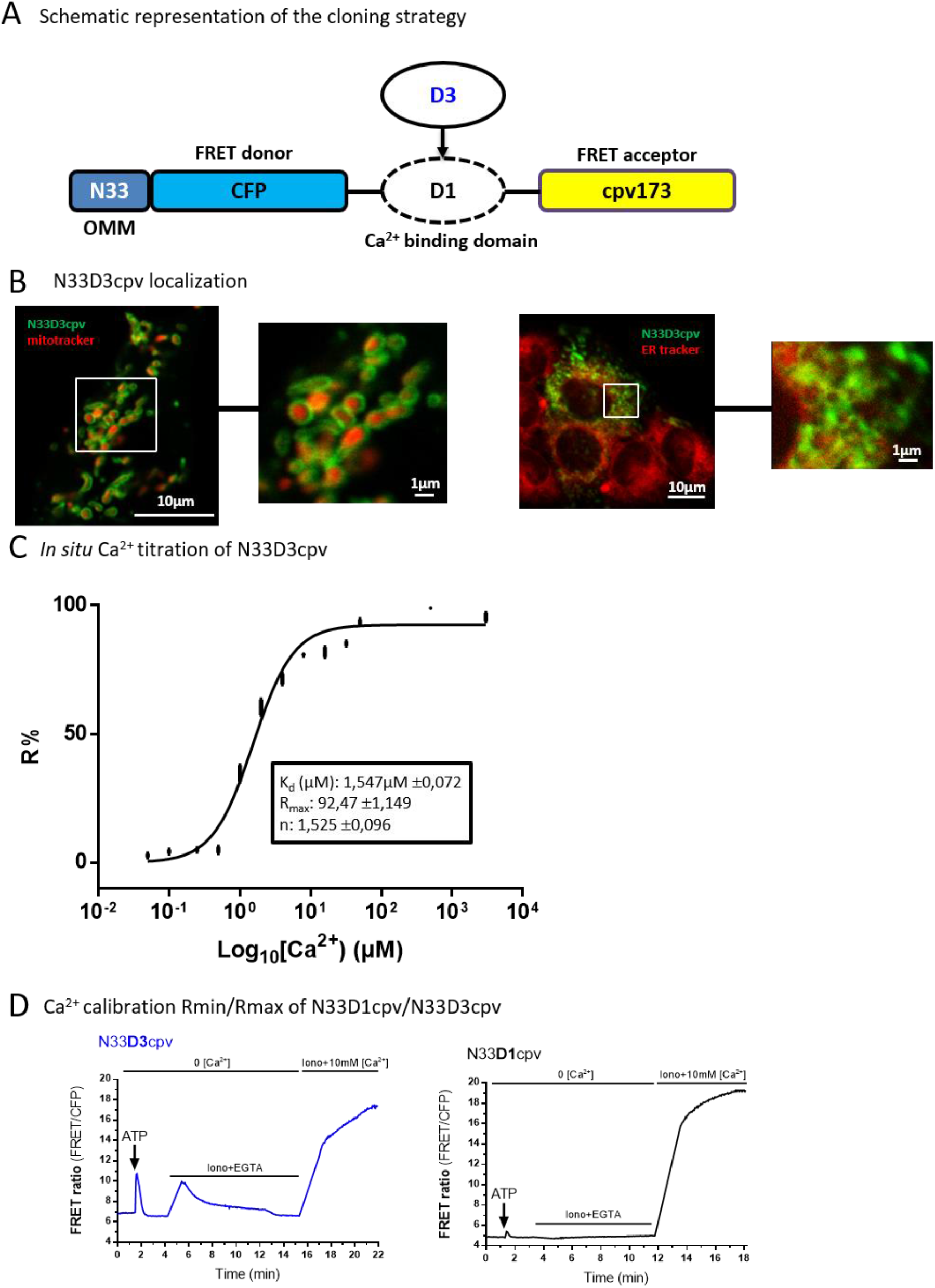
Characterization of N33D3cpv. (**A**) Schematic representation of the cloning strategy. Original Ca^2+^ biosensor N33D1cpv is composed of a signal addressing sequence (N33) coding for an outer mitochondrial membrane (OMM) peptide, the FRET donor (CFP: Cyan Fluorescent Protein), the Ca^2+^-binding domain D1 and the FRET acceptor (cpv173: circularly permutated venus protein). N33D3cpv was generated by replacing D1 Ca^2+^-binding domain with the D3 domain. (**B**) (left panel) Confocal images of H9c2 cell expressing N33D3cpv biosensor (green) and stained with a mitochondrial marker (mitotracker deep red). (right panel) Confocal images of H9c2 cell expressing N33D3cpv biosensor (green) and stained with an ER marker (ER tracker red). Zoomed-in panels for each confocal image are represented on the original image by a white square (**C**) *In situ* Ca^2+^ titration assay of N33D3cpv with the fit values shown in the box. Data plotted: mean ± SEM (n ≥ 9) cells for each [Ca^2+^]. (D) Representative kinetics of FRET ratio (FRET/CFP) of H9C2 cells stimulated with 100μM ATP in Ca^2+^ free extracellular medium then permeabilized with ionomycin (5μM) in an intracellular medium containing EGTA (600μM) and BAPTA-AM (5μM) thenfinally perfused with an intracellular medium containing CaCl_2_ (10mM). (Left panel) N33D3cpv. (Right panel) N33D1cpv. Raw values of FRET ratio are presented (FRET canal/CFP).

### Ca^2+^ affinity of the new generated N33D3cpv indicator

A better sensitivity to Ca^2+^ must enable the detection of low Ca^2+^ levels but it should also modify the kinetics of detection. To confirm this point, we used three different Ca^2+^-mobilizing stimuli and compared the responses measured by N33D1cpv and N33D3cpv indicators. We have chosen fast, slow and long-lasting kinetics of Ca^2+^ release on the surface of the OMM induced by ATP or oxygen glucose deprivation (OGD), respectively without external Ca^2+^. We also used cyclopiazonic acid (CPA) to block SERCA pump activity in order to visualize the slow ER Ca^2+^ leakage. To compare both indicators’ sensitivities and dynamics, we plotted them on the same graph by normalizing the FRET-ratio (F) with the baseline FRET-ratio value (F_0_). At first, we compared the amplitude of the Ca^2+^ response following an ATP stimulus and we observed that the peak of averaged F/F_0_ ratio was greater with N33D3cpv than with N33D1cpv: 1.573 and 1.107, respectively (Figure 2A). Second, we stimulated with CPA, a SERCA pump blocker that uncovers the slow passive Ca^2+^ leakage from the endoplasmic reticulum (ER). The peak of averaged F/F_0_ ratio was greater with N33D3cpv than with N33D1cpv, 1.426 and 1.123 respectively and we observed also a difference in the decay of Ca^2+^ level (Figure 2B). Third, we performed an OGD to compare the indicators’ responses and we observed that the peak of averaged F/F_0_ ratio was once again greater with N33D3cpv than with N33D1cpv, 1.439 and 1.131 respectively (Figure 2C). As expected, due to its higher Kd for Ca^2+^ (Kds at 18,61μM and 135,41μM), N33D1cpv biosensor was not sensitive enough to efficiently discriminate the variations in [Ca^2+^]_OMM_ in these three conditions (Figure 2D). Indeed, after calibration, [Ca^2+^]_OMM_ estimated with N33D1cpv biosensor, in resting or stimulated (ATP, CPA or OGD) H9c2 rat cardiomyoblasts, showed a lot of negative values that reported a measure below the dynamic range of the biosensor (Figure 2D, ΔR% between 0,2-6). Conversely, N33D3cpv allowed us to perform accurate and reproducible measurements of [Ca^2+^]_OMM_ of 0.142±0.108 μM, 0,442±0.164 μM 0.426±0.082 μM, 0.434±0.051 μM, 0,2521±0.080 μM and 0.474±0.040 μM in basal, 50μM ATP, 100μM ATP, 5μM CPA, 20μM CPA and OGD conditions, respectively (Supplemental Figure E and Figure 2E, ΔR% between 17-40). Altogether, these results clearly demonstrate the enhanced sensitivity of the new N33D3cpv sensor and its ability to detect lower variations in [Ca^2+^]_OMM_.

**Figure 2.**
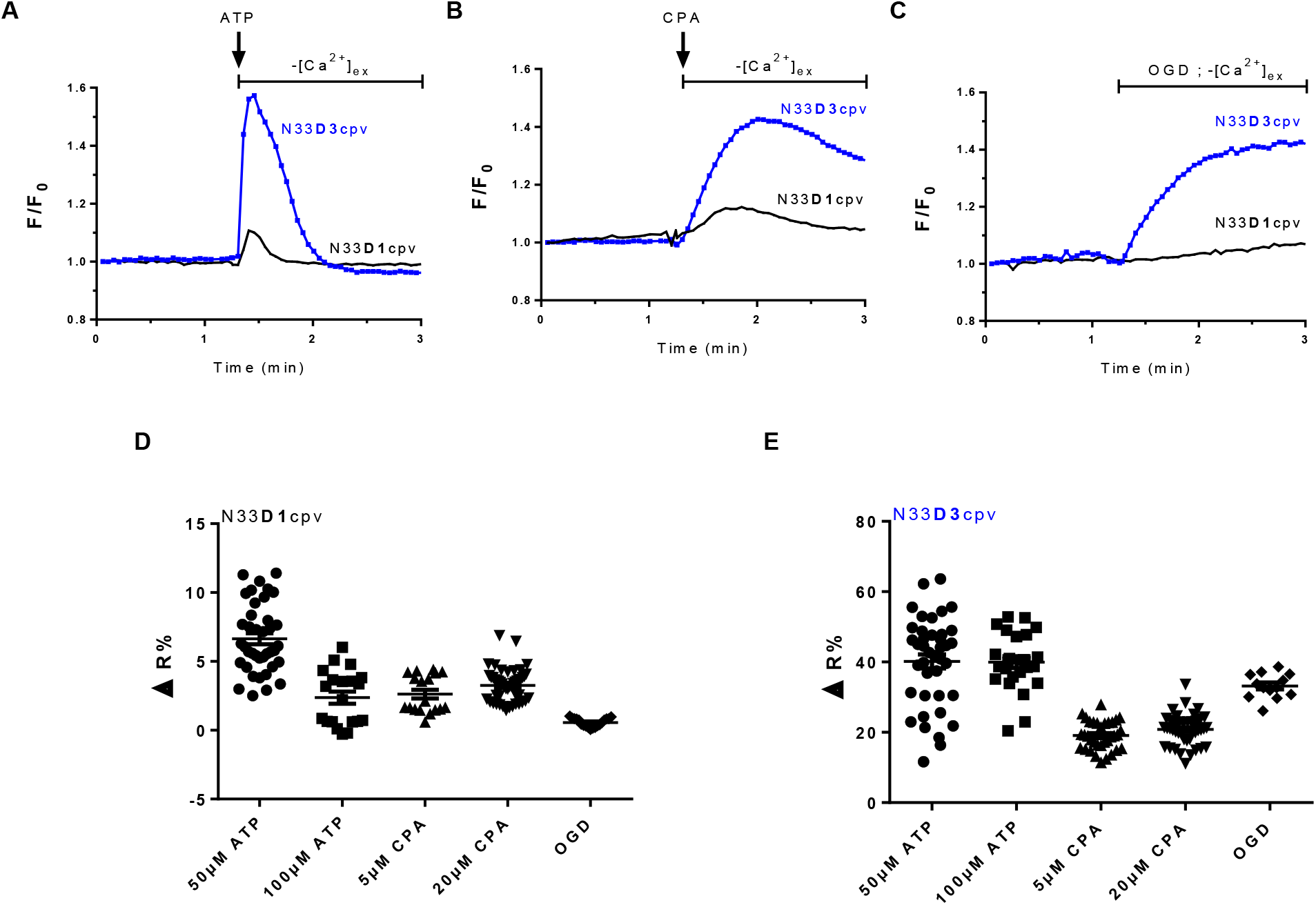
Sensitivity of N33D1cpv and N33D3cpv biosensors in OMM. (**A**) Change in [Ca^2+^]_OMM_ induced by a 100 μM ATP stimulation, using either N33D1cpv (black) or N33D3cpv (blue) in absence of external Ca^2+^. (**B**) Changes in [Ca^2+^]_OMM_ induced by 5 μM cyclopiazonic acid (CPA) stimulation, using N33D1cpv (black) or N33D3cpv (blue) in absence of external Ca^2+^. (**C**) Changes in [Ca^2+^]_OMM_ occurring during an oxygen glucose deprivation (OGD), using N33D1cpv (black) or N33D3cpv (blue) in absence of external Ca^2+^. (**A-C**) Representative average FRET-ratio (F) normalized with the baseline FRET-ratio value (F_0_). (**D-E**) Dot plots represent the mean ±SEM of ΔR% of the 2 probes (N33D1cpv and N33D3cpv, respectively), where ΔR% is calculated as % of the steady-state value (R_1_) and its maximum value (R_2_) after drugs stimulation (ATP, CPA) or OGD. N=3-4, Figure 2D n=39, 20, 18, 48, 19 and Figure 2E n=40, 25, 34, 45, 13 respectively.

### Physiological application of the newly generated N33D3cpv Ca^2+^ indicator

In ischemic condition, the lack of energy supply ceases Ca^2+^ ATPase pumps and depletes ER Ca^2+^ stores. Indeed, we previously demonstrated the rapid decrease of cytosolic and mitochondrial ATP levels upon ischemia. This decrease was concomitant with a rapid release of Ca^2+^ in the cytosol and in the mitochondria (Gouriou et al., 2020). Nevertheless, this mechanism of ER Ca^2+^ depletion remains unclear. Despite the fact that IP3Rs are the major release Ca^2+^ channels in non-excitable cells, their contribution during the ischemic period has not been assessed. We used a model of types-I/II/III IP3Rs triple knock-out (TKO) (Ando et al., 2018) to study their contribution in hypoxia-induced Ca^2+^ leak by using our new N33D3cpv indicator.

First, we controlled the TKO IP3Rs HeLa cells model. Immunoblotting against IP3R1 isoform allowed us to show that, unlike the WT samples, no band at the expected size of about 314kDa was detected in the TKO samples (Supplemental Figure A). A functional assay was done through the measurement of the Ca^2+^ levels using N33D3cpv indicator on stimulated HeLa cells with ATP. We observed an IP3-mediated Ca^2+^ release in WT cells and no response in TKO cells (Supplemental Figure B). We thus confirmed the absence of IP3Rs activity in this model of TKO HeLa cells.

With the objective to assess the implication of IP3Rs in ER Ca^2+^ leak, we assessed the native activity of IP3R while blocking SERCA activity with CPA. We compared the Ca^2+^ responses induced by CPA in WT and TKO HeLa cells with D3cpv (Figure 3A) and N33D3cpv indicators (Figure 3B). As reported in figure 3, D3cpv was unable to detect any difference in the cytosol whereas N33D3cpv biosensor detected a significant decrease in the slope of Ca^2+^ accumulation around mitochondria in TKO IP3Rs (0.101±0.038) as compared to WT HeLa cells (0.154±0.052). Although these results may seem paradoxical, they are in the same line of evidence with prior studies which could not report difference in [Ca^2+^]_cyto_ after blockage of SERCA pumps in TKO and WT cells; whatever the biosensor used: Fura-2 (Kd 0.220μM)(Ando et al., 2018), GEM-GECO (Kd 0.340μM)(Yue et al., 2020). These prior publications mostly supported the fact that TKO had no change in their ER Ca^2+^ stores. Interestingly, by means of GEM-CEPIA1er (Kd 558μM) or R-CEPIA1er (Kd 565μM) (Yue et al., 2020), Yue et al reported a small decrease in the rate of ER Ca^2+^ release, what could be explained by the decrease in ER Ca^2+^ leak, that we observed on the surface of the OMM (Fig. 3C). IP3R1 has been previously reported to exert an endogenous activity as leak channel in non-stimulated cells (Bandara et al., 2013; Kasri et al., 2006). The Ca^2+^ decrease that we observed on the surface of the OMM could thus be related to the loss of native IP3R activity or an increased Ca^2+^ removal by pumps or transporters (so as MCU in mitochondria). The latter hypothesis is very unlikely since SERCA pumps are inhibited by CPA in our experiments. In addition, Yue et al additionally reported a decrease in SERCA-mediated Ca^2+^ uptake in TKO cells that would have been expected to enhance Ca^2+^ accumulation on the surface of the OMM. Finally, the increase in MAMs width, previously reported in the TKO cells(Bartok et al., 2019), is expected to decrease MCU-mediated Ca^2+^ uptake in mitochondria, that would have also led to an increase in Ca^2+^ accumulation on the surface of the OMM. Altogether these results rather validate the hypothesis that endogenous IP3R activity participates in the ER Ca^2+^ leak when SERCA activity is pharmacologically inhibited.

**Figure 3.**
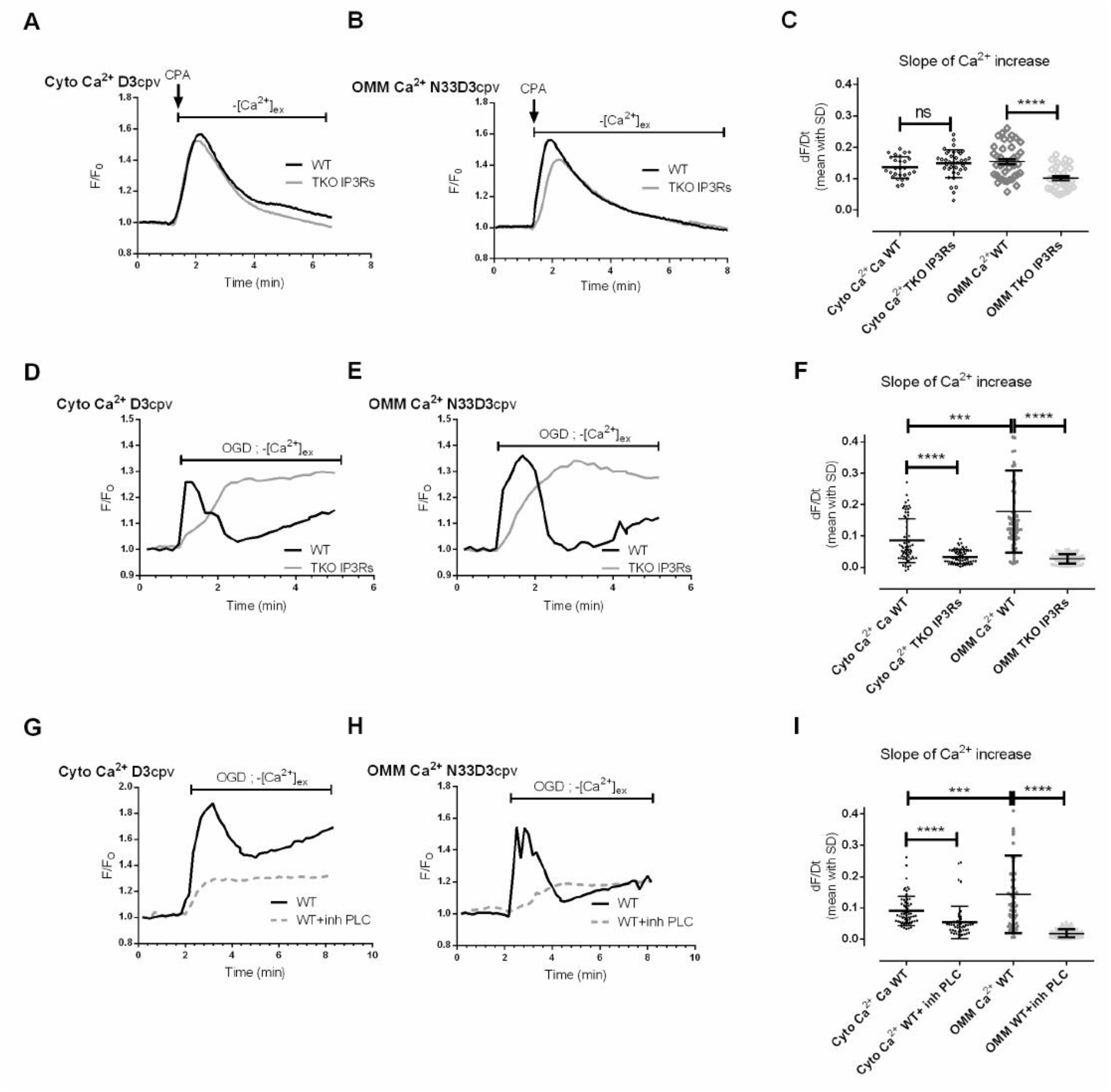
N33D3cpv Ca^2+^ biosensor study the role of IP3R channels in the passive ER Ca2+ leak. Time trace shows [Ca^2+^]_cyto_ (**A**) or [Ca^2+^]_OMM_ (**B**) measured with D3cpv and N33D3cpv, respectively, in absence of external Ca^2+^, in WT (black) and TKO IP3Rs (grey) HeLa cells. A 5 μM cyclopiazonic acid (CPA) stimulation was applied to block SERCA activity in order to reveal the ER Ca^2+^ leak. Representative average FRET-ratio (F) normalized with the baseline FRET-ratio value (F_0_). (**C**) Slope of the Ca^2+^ increase induced by cyclopiazonic acid (CPA) stimulation in WT and TKO IP3Rs HeLa cells N=3-4 n=27, 37, 42, 32 respectively. Time trace shows [Ca^2+^]_cyto_ (**D**) or [Ca^2+^]_OMM_ (**E**) measured with D3cpv and N33D3cpv, respectively, in absence of external Ca^2+^, in WT (black) and TKO IP3Rs (grey) HeLa cells. These cells were subjected to an oxygen glucose deprivation (OGD). Representative average FRET-ratio (F) normalized with the baseline FRET-ratio value (F_0_). (**F**) Slope of the Ca^2+^ increase induced by OGD in WT and TKO IP3Rs HeLa cells. N=3 n=67, 64, 75, 83, respectively. Time trace shows [Ca^2+^]_cyto_ (**G**) or [Ca^2+^]_OMM_ (**H**) measured with D3cpv and N33D3cpv, respectively, in absence of external Ca^2+^, in WT (black) and TKO IP3Rs (grey) HeLa cells. HeLa cells expressing D3cpv or N33D3cpv were treated with 10μM U73122 during an oxygen glucose deprivation (OGD). (I) Slope of the Ca^2+^ increase induced by OGD in WT HeLa cells with or without PLC inhibitor (10μM U73122). N=3 n=65, 50, 57, 84, respectively. Representative average FRET-ratio (F) normalized with the baseline FRET-ratio value (F_0_). (**C, F, I**) Data shown represent the mean with standard deviation (SD) of at least 3-4 independent experiments (ns p≥0.05, * p<0.05, ** p<0.01, *** p<0.001, **** p<0.0001).

We thus wondered whether the native activity of IP3Rs contributed to the ER Ca^2+^ leak during hypoxia. When Hela cells were incubated under oxygen glucose deprivation (OGD), an ER Ca^2+^ leak occurred which could be measured indirectly through the sustained Ca^2+^ increase both in the cytosol, [Ca^2+^]_cyto_ (Figure 3D) and in OMM surface, [Ca^2+^]_OMM_ (Figure 3E). In the supplemental figures C&D, we reported that the precision of the measurements of steady-state [Ca^2+^] by D3cpv (SD: 0.097 and 0.102 for WT and TKO; Supp Fig. C) was 2-fold below the one of N33D3cpv (SD: 0.053 and 0.055 for WT and TKO; Supp Fig. D). Consequently, N33D3cpv could detect a smaller variation in [Ca^2+^] in WT than in TKO HeLa cells, at 0.098 ±0.064 μM and 0.120 ±0.068 μM, respectively (Supp Fig. C&D). We then analyzed the slope of the rise in [Ca^2+^]_cyto_ and in [Ca^2+^]_OMM_ and we found a significantly slower Ca^2+^ increase rate in TKO compared to WT HeLa cells both in the cytosol and on the OMM (Figure 3F). Interestingly, we detected a faster Ca^2+^ increase at the OMM level compared to the cytosol in WT cells but no difference in TKO cells. This result suggests a greater activity of IP3R on the surface of the OMM than in the whole ER membrane (Bartok et al., 2019). However, IP3R clusters participate as physical tethers in MAMs and the loss of IP3Rs has been reported to decrease the frequency of tight contact sites (10-50nm) in MAMs of TKO cells. This modification in MAMs structure could in turn delay the rise of Ca^2+^ on the surface of the OMM. In order to determine to which extent (i) the loss of IP3Rs activity and (ii) the modification in ER-mitochondrial contacts were involved in the observed decrease in the rate of Ca^2+^ accumulation on the surface of the OMMs during the hypoxia, we incubated WT Hela cells with U73122 to inhibit phospholipase C (PLC), which has a crucial role in the initiation of the activation of IP3Rs (Berridge, 1987). WT Hela cells were incubated with 10 μM U73122 (Figure 3G, 3H) and we detected a reduced rate of Ca^2+^ increase in the cytosol and in OMM surface in WT+ PLC inhibitor compared to WT HeLa (Figure 3I). Altogether, these results confirmed that an endogenous IP3R activity participated in the passive ER Ca^2+^ leak occurring during hypoxia and was more specifically localized in the MAMs.

In conclusion, thanks to our newly designed D3cpv biosensor addressed to OMM, we showed that the endogenous IP3R activity participates in ER Ca^2+^ leak during hypoxia. This N33D3cpv biosensor allows very sensitive Ca^2+^ measurement on the surface of mitochondria and will help further researches in the field of ER-mitochondria homeostasis.

## Material and Methods

### N33D3cpv construct strategy

A 1998 bp BamHI-XhoI fragment encompassing D3cpv was prepared from pcDNA-D3cpv, gifted from Roger Tsien (Addgene plasmid # 36323)(Palmer et al., 2006), and subcloned between BamHI and XhoI restriction sites in pcDNA-N33D1cpv, gifted from Tullio Pozzan(Giacomello et al., 2010), to generate pcDNA-N33D3cpv (7583 bp). Created plasmid DNA sequence was confirmed by Sanger sequencing.

### Cell culture and transfection

Rat cardiomyoblasts H9c2 (ATCC, CRL-1446) and WT / TKO -HeLa cells (kindly provided by Katsuhiko Mikoshiba at SIAIS, ShanghaiTech University, Shanghai, China) were routinely cultured in Dulbecco’s Modified Eagle Medium (Gibco) supplemented with 10% fetal calf serum (PAN-biotech), 100 U/mL penicillin (Gibco), 100 μg/mL streptomycin (Gibco), 2 mM L-glutamine (Gibco) and incubated at 37°C in 5% CO_2_ in a damp atmosphere. Cells were regularly passaged by single-cell dissociation with 0.05% trypsin-EDTA (Gibco). For generation of transient transfectants, all DNAs (N33D1cpv and N33D3cpv) were transfected into cells using DharmaFECT Duo (Dharmacon, T-2010-03) according to manufacturer’s instructions. Cells were plated on glass coverslips 24 hours before transfection and experiments were performed 48 hours after.

### Immunoblotting

For Western-blotting, cell lysates were obtained by treating the cell monolayer with RIPA buffer complemented with protease and phosphatase inhibitors. Protein lysates were then cleared by centrifugation (17’000g for 20min). Total protein concentration was determined using bicinchoninic acid protein assay (Interchim, UP40840) and 25μg of denatured and reduced proteins of each sample was loaded on a 6% SDS-PAGE. After SDS-PAGE migration and electroblotting on polyvinylidene fluoride, the membranes were blocked with 5% non-fat milk and then incubated with the specific primary antibodies [rabbit anti-IP3R1 (Santa-Cruz, sc-28614; 1/500) and rabbit anti-TUBULIN (Santa-Cruz, sc-5286; 1/500)]. Blots were incubated with horseradish peroxidase (HRP)-coupled sheep anti-mouse IgG (GE Healthcare, NA931VS; 1/10000) and (HRP)-coupled goat anti-rabbit IgG (GE Healthcare, NA934VS; 1/10000), and developed with Clarity Western ECL Substrate (BioRad, 1705060). The band intensity was determined using Image Lab software (Bio-Rad).

### Wide-field imaging

Living cells were imaged on an inverted epifluorescence microscope Leica DMi6000B using a 40x oil-immersion objective, with Lambda DG4 wavelength-switch xenon light source (Sutter Instruments), equipped with an ORCA-Flash4.0 digital CMOS camera C11440 (Hamamatsu). Pictures have been acquired with 200 ms acquisition time per frame, 10% fluorescence intensity manager (FIM) and an interval of 1 second for time-lapse. Cameleon fluorescent proteins were excited at a wavelength of 430nm and emissions were collected at 480nm and 530nm. Fluorescence ratio imaging was analysed using MetaFluor software (Molecular Devices). Experiments were carried out in controlled environment at 37°C and cells were placed in Ca^2+^-free buffer containing 140mM NaCl, 5mM KCl, 1mM MgCl_2_, 10mM HEPES and 10 mM Glucose, adjusted to pH 7.4, supplemented with 1 mM EGTA.

Dynamic range method: In H9C2 cells expressing N33D3cpv or N33D1cpv, Rmin and Rmax were obtained upon permeabilization of the cells with 5μM ionomycin and a Ca^2+^-free buffer containing 140mM NaCl, 5mM KCl, 1mM MgCl_2_, 10mM HEPES and 10 mM Glucose, adjusted to pH 7.4, supplemented with 600μM EGTA. The Rmin was achieved by perfusing the cells with this medium containing 600μM EGTA and 5μM BAPTA-AM. The Rmax was achieved by perfusing the cells with a Ca^2+^ buffer containing 140mM NaCl, 5mM KCl, 1mM MgCl_2_, 10mM HEPES and 10 mM Glucose, adjusted to pH 7.4, supplemented with 10mM CaCl2. R% is calculated as R% = (R − R_min_)/(R_max_ − R_min_) × 100.

In situ Ca^2+^ titration assay: H9C2 cells expressing N33D3cpv were permeabilized with 100μM digitonin in a Ca^2+^-free medium containing 600μM EGTA for 60 sec and then washed 3 times with the same medium without digitonin. Cells were then perfused with Ca^2+^ buffer containing 140mM NaCl, 5mM KCl, 1mM MgCl_2_, 10mM HEPES and 10 mM Glucose, adjusted to pH 7.4, and known Ca^2+^ concentrations. At the end of each experiment, a saturating Ca^2+^ concentration (10mM) was applied. For [Ca^2+^] lower than 0,5μM, the buffer was supplied with BAPTA free acid and Ca^2+^. The free [Ca^2+^] was estimated using MaxChelator. The results obtained were plotted as log_10_[Ca^2+^] (x-axis) and R% (y-axis) and fitted using Prism 9.0 (GraphPad) with the following equation: y = (Rmax1 × x^n1^)/(kd1^n1^ + x^n1^)+(Rmax2 × x^n2^)/(kd2^n2^ + x^n2^).

### Oxygen-Glucose Deprivation (OGD) experiments

Cells were washed twice, placed in Ca^2+^-containing buffer having 140mM NaCl, 5mM KCl, 1mM MgCl_2_, 10 mM HEPES and 2mM Na_2_S_2_O_4_, adjusted to pH 7.4, supplemented with 2mM CaCl_2_. Cells were placed in a specifically manufactured bio-incubator (NewBrunswik, Galaxy 48R) connected with a 100% N_2_ bottle. Oxygen level and temperature were monitored at 0.5% and at 37°C respectively. An Okolab system with a specific hypoxic chamber was used to control the environmental constants (temperature, humidity and oxygen levels).

### Confocal imaging

Living cells were imaged on an inverted confocal microscope Nikon A1R+ system using 40x oil-immersion objective with Argon laser (488-514nm) for the Cameleon excitation, Diode (642nm) for the Mitotracker Deep Red and diode (560nm) for the ERtracker Red. The images were acquired on living cells plated on glass coverslips.

### Statistical analysis

Data processing and statistical analyses were conducted with Prism 9.0 (GraphPad) software. Before proceeding to any analysis, the normality of the samples was evaluated. Unpaired *t*-test (for normal distribution) or Mann–Whitney test (for non-normal distribution) was used unless stated otherwise in the figure legends. *p*-values are indicated in figures. Data show mean with standard deviations (SD) calculated from at least three independent experiments. For single-cell imaging analysis, statistics were performed on *n* = number of cells to assess single-cell effect as well as heterogeneity between them. Both *N* and *n* values are indicated in the figure legends. A *p* value < 0.05 was considered significant.

## Acknowledgments

We would like to thank Roger Tsien for the D3cpv construct (University of California, San Diego, CA) and Marta Giacomelo, Paola Pizzo and Tullio Pozzan for the N33D1cpv construct (Department of Biomedical Sciences, University of Padova, Italy). HeLa WT and TKO IP3Rs cells were a gift from Katsuhiko Mikoshiba, SIAIS, ShanghaiTech University, Shanghai, China. We address a special thanks to Mélanie Paillard and Mariam Wehbi for proofreading the manuscript.

