## Supplemental figure for "High-sensitivity calcium biosensor on the mitochondrial surface reveals that IP3R channels participate in the reticular Ca^2+^ leak towards mitochondria"

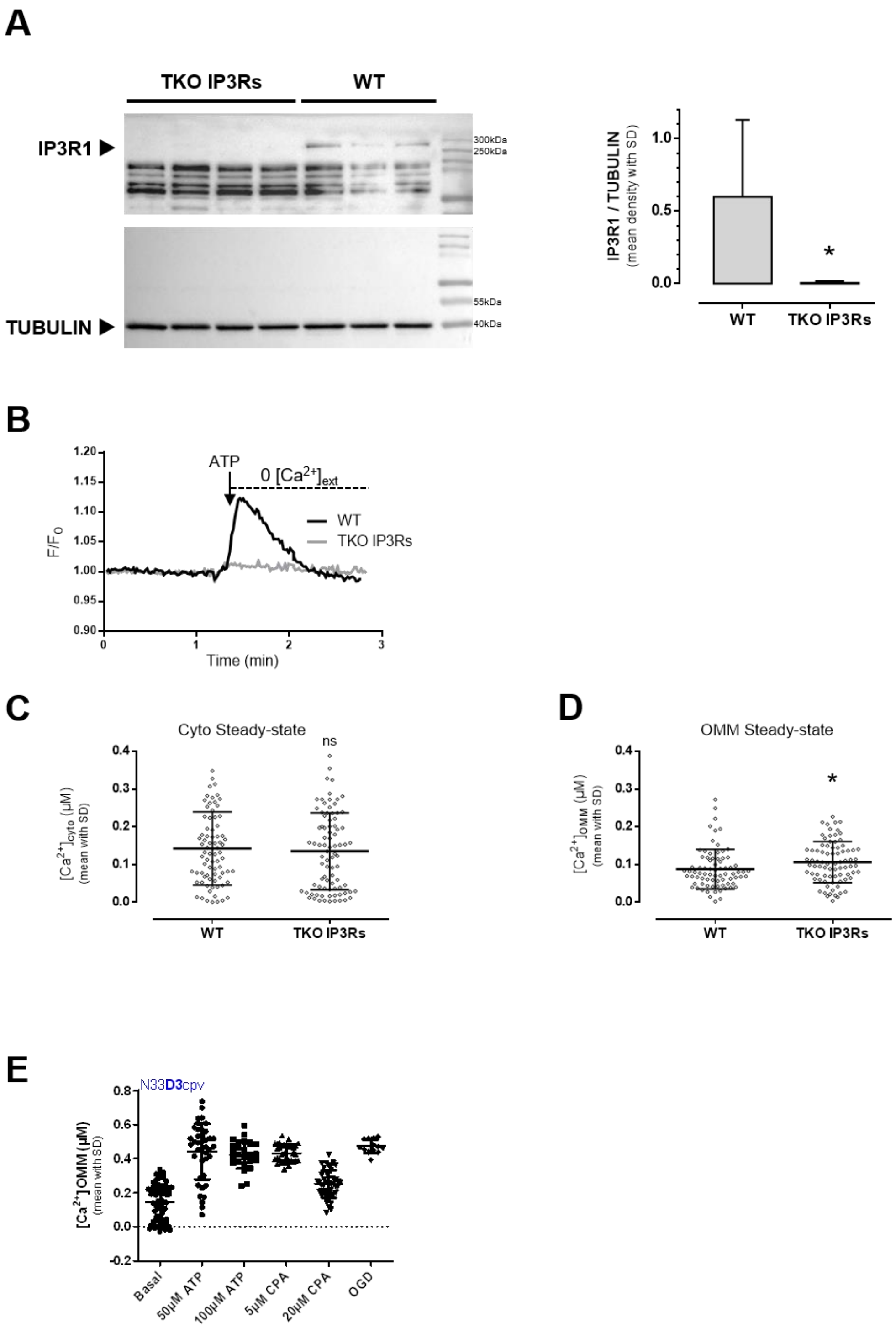


***Supplementary figure. D3cpv $Ca^{2+}$ biosensor to study the role of IP3R channels in the passive ER Ca2+ leak***

(A) Immunoblotting against IP3R1 isoform and tubulin in WT and TKO IP3Rs HeLa cells. Data shown represent the mean with standard deviation (SD) of 3 independent experiments, (* $p<0.05$). (B) $[Ca^{2+}]_{OMM}$ was estimated using N33D3cpv biosensor in WT and TKO IP3Rs HeLa cells treated with 100 µM Na, in absence of external $Ca^{2+}$. Representative average FRET-ratio (F) normalized with the baseline FRET-ratio value ($F_0$). (C) Steady-state $[Ca^{2+}]_{cyto}$ in WT and TKO IP3Rs HeLa cells. (D) Steady-state $[Ca^{2+}]_{OMM}$ in WT and TKO IP3Rs HeLa cells. (E) $[Ca^{2+}]_{OMM}$ steady state (basal) and peak measurements upon ATP, CPA and OGD treatment protocol with the N33D3cpv sensor.
